# The fitness cost of a congenital heart defect shapes its genetic architecture

**DOI:** 10.1101/531988

**Authors:** Ehiole Akhirome, Suk D. Regmi, Rachel A. Magnan, Nelson Ugwu, Yidan Qin, Claire E. Schulkey, James M. Cheverud, Patrick Y. Jay

## Abstract

**Background:** In newborns, severe congenital heart defects are rarer than mild ones. The reason why is unknown, but presumably related to a liability threshold that rises with the severity of a defect. Because the same genetic mutation can cause different defects, other variables may contribute to pushing an individual past a defect-specific liability threshold. We consider here how variables in the genetic architecture of a heart defect depend upon its fitness cost, as defined by the likelihood of survival to reproductive age in natural history studies.

**Methods:** We phenotyped ~10,000 *Nkx2-5*^+/-^ newborn mice, a model of human congenital heart disease, from two inbred strain crosses. Genome-wide association analyses detected loci that modify the risk of an atrial septal defect, membranous or muscular ventricular septal defect, or atrioventricular septal defect. The number of loci, heritability and quantitative effects on risk of pairwise (G×G_*Nkx*_) and higher-order (G×G×G_*Nkx*_) epistasis between the loci and *Nkx2-5* mutation were examined as a function of the fitness cost of a defect.

**Results:** *Nkx2-5*^+/-^ mice have pleiotropic heart defects; about 70% have normal hearts. The model recapitulates the epidemiological relationship between the severity and incidence of a heart defect. Neither the number of modifier loci nor heritability depends upon the severity of a defect, but G×G_*Nkx*_ and G×G×G_*Nkx*_ effects on risk do. Interestingly, G×G×G_*Nkx*_ effects are three times more likely to suppress risk when the genotypes at the first two loci are homozygous and from the same, rather than opposite strains in a cross. Syn- and anti-homozygous genotypes at G×G×G_*Nkx*_ interactions can have an especially large impact on the risk of an atrioventricular septal defect.

**Conclusions:** Given a modestly penetrant mutation, epistasis contributes more to the risk of severe than mild congenital heart defect. Conversely, genetic compatibility between interacting genes, as indicated by the protective effects of syn-homozygosity at G×G×G_*Nkx*_ interactions, plays a newfound role in the robustness of cardiac development. The experimental model offers practical insights into the nature of genetic risk in congenital heart disease. The results more fundamentally address a longstanding question regarding how mutational robustness could arise from natural selection.

## INTRODUCTION

Congenital heart disease (CHD) is recognized as the most common birth defect, but it actually encompasses multiple, anatomically distinct phenotypes in which the severity and incidence of a defect are inversely related. For example, secundum atrial septal defects (ASD), which are compatible with survival into middle age, are three times more common than atrioventricular septal defects (AVSD), which can cause death during infancy.^1, 2^ There is no simple explanation for the epidemiological relationship, such as a genotype-phenotype relationship or lower frequency of the causes of severe defects. The relationship does imply that each defect has a liability threshold related to its severity. The higher the threshold, the rarer a defect is. The developmental pathways that lead to a severe defect appear more robust to perturbation than the ones to a mild defect.

Questions regarding how the fitness cost of a defect relates to the robustness of the underlying pathways are practical and fundamental. An appreciation of the genetic architecture of a heart defect – i.e., the genes involved, alleles, interactions, and their effects – should inform genetic risk models and potential strategies for prevention. The genetic architecture clearly encompasses more than the proximate cause of disease. As a rule, the same deleterious mutation can manifest as a mild, moderate or severe defect in different individuals. Modifier genes affect the phenotypic outcome.^3^ Genetic heterogeneity and rare alleles unfortunately hamper the detection of epistasis in humans, which in turn obscures the elucidation of complex genetic principles. Inbred strain crosses in mutant mouse models can circumvent these limitations. The number of alleles at a locus can be limited to two, and the frequencies of both alleles can be kept equally common.

Through inbred strain crosses of *Nkx2-5*^+/-^ mice, a model of non-syndromic human CHD,^4, 5^ we have shown that the incidence of specific defects can vary between genetic backgrounds. Genetic interactions modulate the susceptibility of cardiac developmental pathways to the deleterious mutation.^6, 7^ In addition, first-generation (F1) hybrids of the inbred strains C57BL/6N and FVB/N or A/J have a lower incidence of congenital heart defects than *Nkx2-5*^+/-^ mice in the C57BL/6N background or the second-generation (F2) offspring of backcrosses to the parental strains or intercrosses (F1×F1). The observations suggest hybrid vigor or heterosis. That is, genome-wide heterozygosity increases the average level of robustness in a population.^6^.

The observations also imply that variation in the robustness to the *Nkx2-5* mutation results from variation in the effects of genetic interactions. Epistasis contributes quantifiably to fitness traits in simple organisms, such as yeast,^8^ *Drosophila*^9^ and *C. elegans*.^10^ On the other hand, statistical genetic evidence for a contribution of epistasis to phenotypes in higher organisms is scant, and theoretical analyses arrive at disparate conclusions regarding the significance of epistasis in complex traits and disease.^11, 12^ Why the results conflict is hotly debated.^13^ One, uncommonly discussed reason is that the contribution of epistasis could vary with the fitness cost of a phenotype.^12^ Natural selection indisputably eliminates deleterious mutations, but selection may also increase the robustness of pathways to perturbation by shaping genetic interactions and networks.^14^

We reasoned that a comparative analysis of the genetic architecture of four CHD phenotypes under the same mutation could illuminate the problem. The four defects – ASDs, muscular and membranous ventricular septal defects (VSD), and AVSD – have mild to severe fitness costs, as defined by the likelihood of survival to reproductive age in humans.^2^ We determined the number and effects of quantitative trait loci (QTLs) on the risk of each heart defect in *Nkx2-5*^+/-^ mice from two different inbred strain crosses. Each QTL represents a pairwise interaction between a modifier gene and *Nkx2-5* (G×G_*Nkx*_). In addition, we examined the non-linear effects of higher-order interactions between two QTLs and *Nkx2-5* (G×G×G_*Nkx*_). Empirical data are sorely needed to understand the nature of genetic risk in CHD. More fundamentally, a quantitative analysis of epistasis in a set of phenotypes that vary in fitness costs could inform a longstanding debate regarding the origins of mutational robustness.^15^

## RESULTS

We phenotyped ~10,000 *Nkx2-5*^+/-^ newborns from intercrosses between the inbred strains A/J and C57BL/6N (A×B, N = 2999) and FVB/N and C57BL/6N (F×B; N = 6958). An ASD is the most common defect associated with human *NKX2-5* mutation.^5, 16^ An ASD is also most common in the mouse crosses, followed by membranous and muscular ventricular septal defects (VSD) and AVSD (Fig. 1a). The two anatomic types of VSD have different developmental bases, but similar pathological consequences. We note this because epidemiological studies typically combine membranous and muscular VSDs, making VSD appear more common than ASD.^1^ Both intercrosses recapitulate the inverse relationship between the severity and incidence of a heart defect (Fig. 1b). The attrition of more severely affected embryos does not explain the distribution of defects because *Nkx2-5*^+/-^ mice are born at the expected Mendelian ratio. In fact, very severe defects, such as double outlet right ventricle and tricuspid atresia, were observed, but there were too few for quantitative genetic analysis. About 70% of *Nkx2-5*^+/-^ mice from either intercross have a normal heart, which indicates the modest penetrance of the mutation.

**Figure 1.**
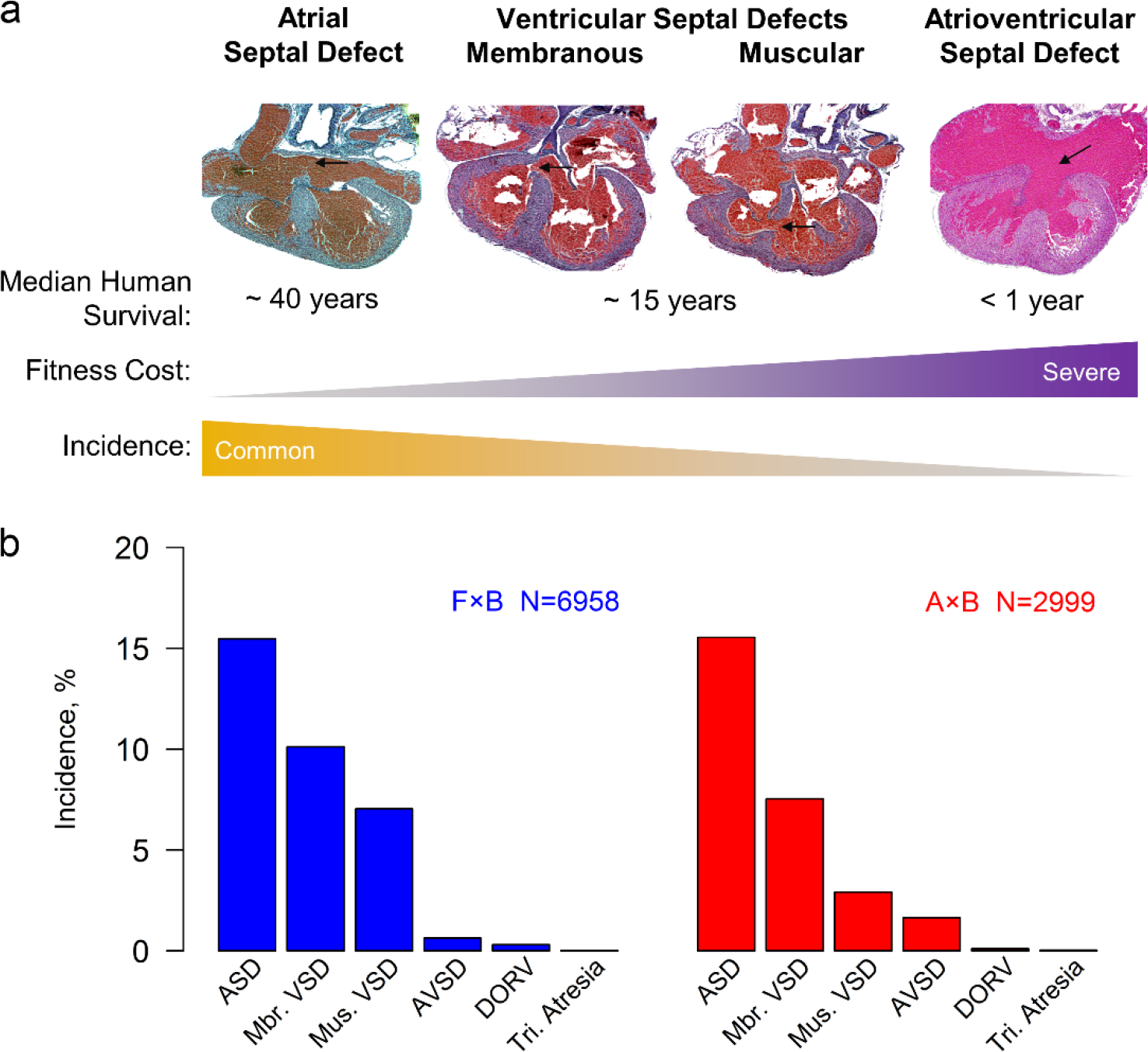
The severity and incidence of a heart defect are negatively correlated. **a,** In humans, mild defects are more common than severe ones. The median survival without surgical intervention defines the severity of a defect ^2^. The panels show representative heart defects from *Nkx2-5*^+/-^ pups. Arrows point to a secundum ASD, membranous and muscular VSDs, and the common atrioventricular canal in an AVSD. **b,** As in humans, the incidences of heart defects in *Nkx2-5*^+/-^ newborns are inversely related with their severity (Kendall’s partial rank correlation τ = −0.845, P < 0.01). Double outlet right ventricle (DORV) was present in 18 and 3 *Nkx2-5*^+/-^ hearts from the F×B and A×B intercrosses respectively. One case of tricuspid atresia was found in each intercross.

The total, quantitative effect of genetic modifiers on risk could be a function of the number of genes, their effect sizes or both. To distinguish these possibilities, we mapped the modifier QTLs in the F2 intercross of A×B and in the combined F2 and advanced intercross generations of F×B (Supplemental Fig. 1). Mapping in an F2 intercross has greater power to detect loci, whereas the combined analysis of an F2 and later generations has greater mapping resolution^17^. We identified QTLs for ASD, membranous and muscular VSD, and AVSD (Fig. 2a, Supplementary Table 1). Each set of QTLs represents G×G_*Nkx*_ interactions in developmental pathways leading to a defect. QTLs that overlap between defects may share a gene involved in the development of multiple cardiac structures.

**Figure 2.**
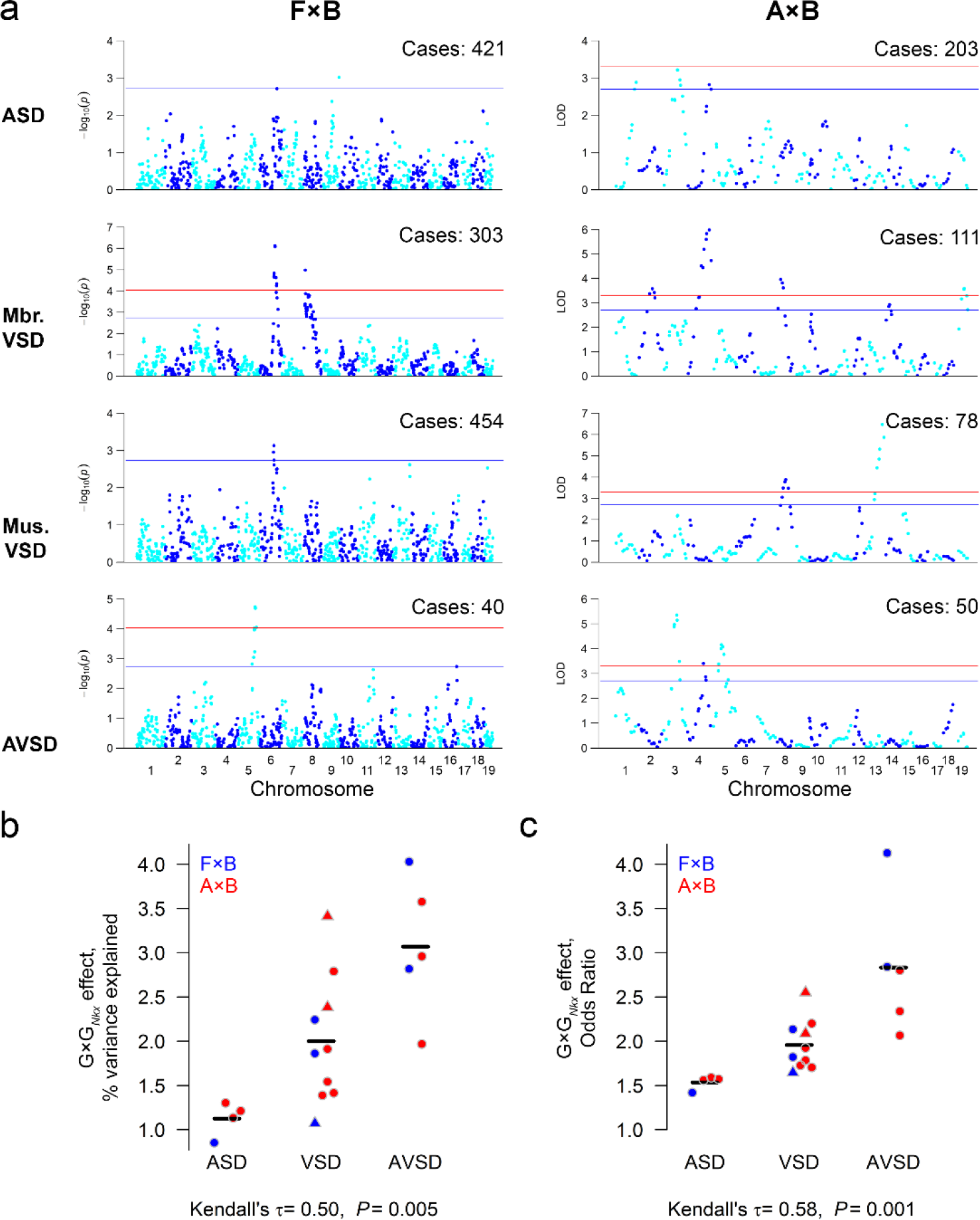
The quantitative effect of G×G_*Nkx*_ interactions on risk correlates with the severity of a heart defect. **a**, Genetic loci modify the risk of specific heart defects in *Nkx2-5*^+/-^ mice from the F×B and A×B intercrosses. The number of affected *Nkx2-5*^+/-^ newborns genotyped for each defect is indicated. Normal *Nkx2-5*^+/-^ littermates were genotyped as controls (N = 330 A×B and 1024 F×B). Suggestive and genome-wide significance thresholds are indicated in blue and red. **b**, **c**, G×G*_Nkx_* effects on risk, quantified as the percent variance explained by the most significant SNP at a locus or the odds ratio for each risk allele, correlate with defect severity. Muscular (▴) and membranous (•) VSD loci are analyzed together because the two VSD types have similar severity. The hatch marks indicate the mean.

The number of detected loci is similar between the four heart defects (Supplemental Fig. 2a). G×G_*Nkx*_ effect sizes, however, vary with the severity of a defect. G×G_*Nkx*_ interactions have a small effect on ASD risk and increasingly larger effects for VSDs and AVSD. The correlation holds whether the effect is calculated as the risk associated with a susceptibility allele or the phenotypic variance explained by a locus (Fig. 2b, c). To account for undetected loci, we estimated the phenotypic variability, i.e., heritability, explained by all genotyped SNPs. The heritability is similar between defects (Supplemental Fig. 2b). The correlation between G×G_*Nkx*_ effect size and severity is not due to a larger total genetic contribution to the risk of severe defects.

While each locus was mapped according to its individual effect on risk, two loci may interact. Representative examples compare the observed to the expected incidences at two-locus genotypes under the null model of non-interacting loci.^18^ Small differences between the observed and expected incidences of ASD and VSD indicate that each locus exerts mainly independent or additive effects (Fig. 3a, b; Supplemental Fig. 3). In contrast, the incidence of AVSD at a two-locus genotype can deviate substantially from the null expectation, indicating a non-linear effect. Some differences are very large relative to the population or the unweighted average incidence of a defect across the nine genotypes (Fig. 3c, d; Supplemental Fig. 3). The latter calculation of incidence permits comparisons between defects of higher-order epistatic effects independently of genotype frequencies. The population and unweighted average incidences are correlated in the inbred strain crosses because there are only two, common alleles per locus. In natural populations, multiple and rare alleles disrupt the correlation and make epistasis difficult to assess.

**Figure 3.**
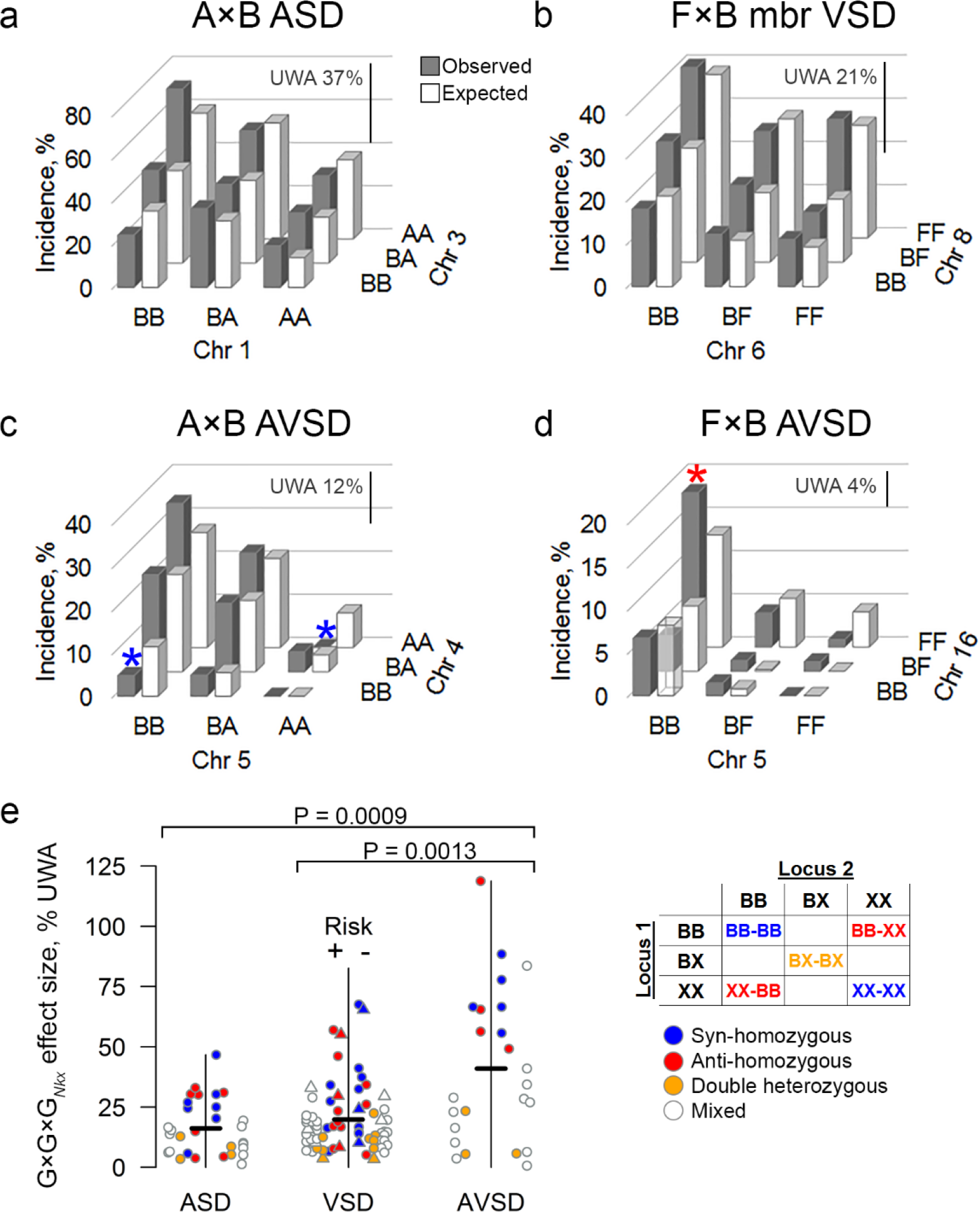
G×G×G_*Nkx*_ effects correlate with the severity of a heart defect. Representative plots depict the observed and expected incidence of a defect at a two-locus genotype. The expected incidence is calculated under the null hypothesis of non-interacting loci. The difference between the observed and expected incidences is the G×G×G_*Nkx*_ effect at a genotype. Not all differences are significant. **a**, **b** ASD and VSD loci mainly contribute to risk independently, as indicated by similar observed and expected incidences. Significant G×G×G_*Nkx*_ effects are small relative to the unweighted average of incidences (UWA, scale bars) across all nine genotypes. **c**, **d**, Epistatic interactions between AVSD loci exert large G×G×G*_Nkx_* effects. Consider two examples (*). The syn-homozygous, two-locus genotypes (AA-AA and BB-BB) in the A×B intercross effectively suppress risk. In contrast, half of the AVSDs in the F×B intercross are associated with an anti-homozygous, Chr 5 BB-Chr 16 FF genotype. **e**, G×G×G*_Nkx_* effect sizes correlate with defect severity (Kendall’s τ = 0.195, P = 0.0017). AVSD effects are larger than ASD and VSD (Mann-Whitney U test). Significant G×G×*G_Nkx_* effects were divided by the UWA and shown as absolute values because the net deviation of observed and expected incidences is zero. The sign indicates whether a significant effect increases (+) or decreases (-) risk. The hatch marks indicate the mean. Effects are color-coded by their two-locus genotypes, as given in the Punnett square for syn- and anti-homozygous, double heterozygous, and mixed genotypes. Alleles: B, C57BL/6N; X, A/J or FVB/N.

Similar to G×*G*_*Nkx*_ interactions, G×G×*G*_*Nkx*_ effect sizes increase with the severity of a defect (Fig. 3e). We calculated the G×G×*G*_*Nkx*_ contribution to the observed incidence of a defect at each of the nine genotypes between every pair of significant and suggestive loci (or the next most significant locus in the two cases where there is just one suggestive locus). Statistically significant G×G×*G*_*Nkx*_ effect sizes vary between defects. AVSD effects are larger than ASD and VSD effects. The relationship is not secondary to an underlying correlation, such as between the observed incidence at a two-locus genotype and the G×G×*G*_*Nkx*_ effect size or the severity of a defect.

Patterns of G×G×*G*_*Nkx*_ effects suggest that positive selection promoted the coadaptation of interacting genes, as opposed to the elimination of intrinsically deleterious alleles. Among significant G×G×*G*_*Nkx*_ effects, two-locus genotypes that are homozygous at both loci for alleles from the same inbred strain or “syn-homozygous”, e.g., BB at both loci, are more likely to lower risk. Conversely, anti-homozygous G×G×*G*_*Nkx*_ effects, e.g., homozygous FF at one locus and BB at the other, are more likely to raise risk. Heterozygosity at either locus is equally likely to raise or lower risk (Fig. 4a). Syn- and anti-homozygous G×G×*G*_*Nkx*_ interactions can have major effects on the risk of a severe defect. For example, one anti-homozygous, two-locus genotype accounts for 50% of all the AVSDs in the F×B intercross (Fig. 3d). The genotypes at either locus are not intrinsically deleterious because *Nkx2-5* wild-type mice, including littermates of affected mutants, do not develop AVSDs. Conversely, both syn-homozygous genotypes at two loci in the A×B intercross effectively suppress AVSD risk (Fig. 3c). In general, no one- or two-locus genotype skews ASD or VSD risk as extremely (Supplemental Figure 3). Internally consistent G×G×*G*_*Nkx*_ effects suggest that selection for coadaptation was a recurrent event in all three inbred strains examined. Protective and deleterious effects of syn- and anti-homozygosity, respectively, are associated with congruent effects at the other syn- and anti-homozygous genotypes at a pair of interacting loci (Fig. 4b-e).

**Figure 4.**
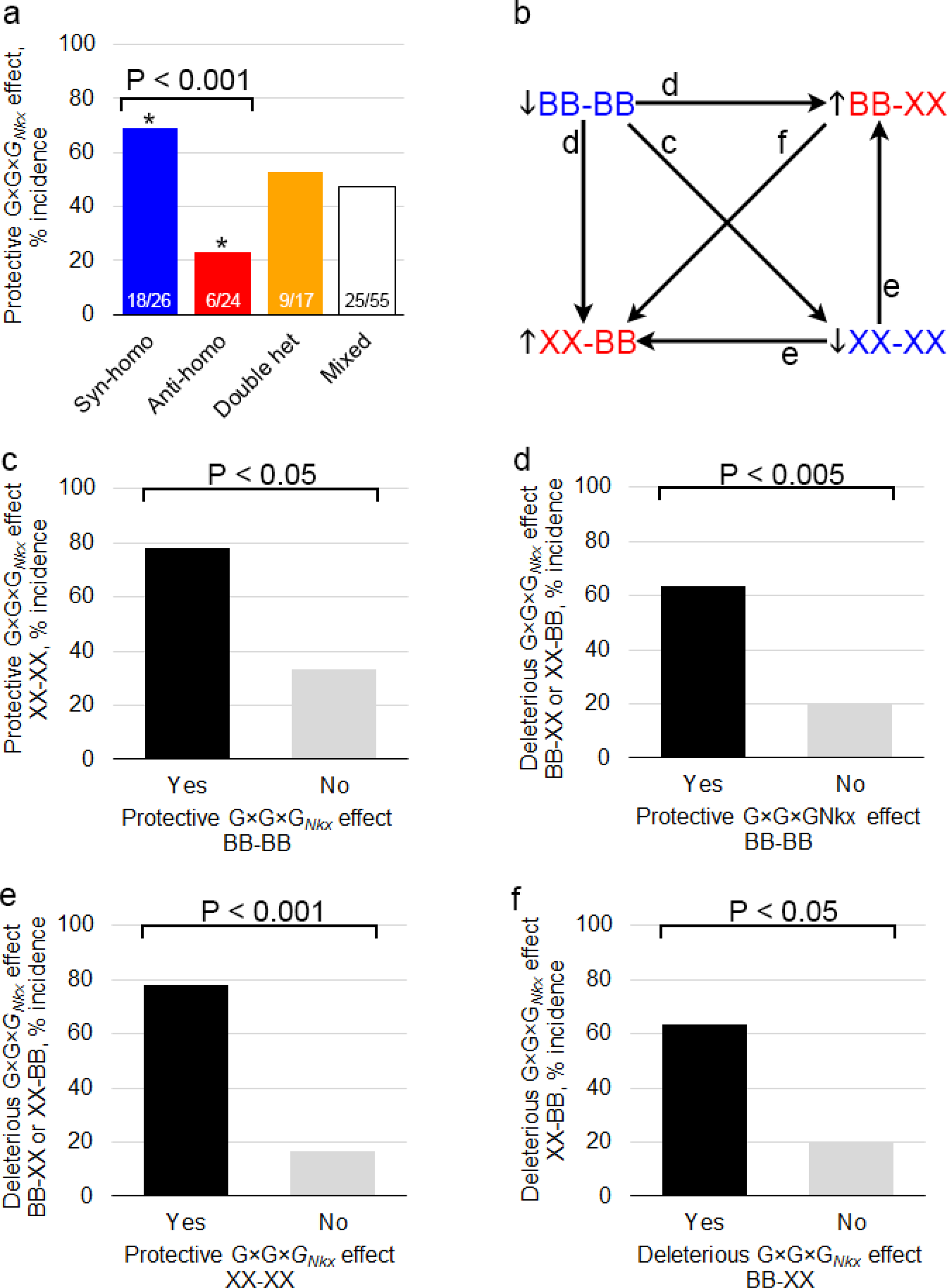
Genetic compatibility between the two loci in a G×G×G*_Nkx_* interaction lowers the risk of heart defects in Nkx2-5^+/-^ mice. a, Statistically significant syn-homozygous G×G×G*_Nkx_* effects between two modifier loci are more likely to be protective than expected by chance or compared to anti-homozygous effects. Anti-homozygous effects are less likely to be protective than expected by chance. Double-heterozygous and mixed genotype effects are equally likely to be protective or deleterious. The fractions of significant effects that are protective are indicated. *, not equal to 50%, P < 0.05. b-f, A significant G×G×G*_Nkx_* effect at a syn-homozygous genotype that is protective (risk ↓) or at an anti-homozygous genotype that is deleterious (risk ↑) is associated with congruent effects at the other syn- and anti-homozygous genotypes. Proportions were compared in two-sided z-tests.

## DISCUSSION

Normal cardiac development is crucial for survival to reproductive age, but CHD phenotypes exact widely ranging costs on fitness. One has to wonder about an evolutionary basis for the fact that severe defects are rarer than mild ones.^19^ Highly penetrant, mostly *de novo* genetic abnormalities account for about one third of all CHD cases^3^, but they have not been associated with the most severe defects. In fact, ASDs accounted for more than 10% of CHD cases in two large, whole-exome sequencing studies that examined the role of *de novo* mutation.^20, 21^ We propose instead that the inverse relationship is a consequence of the genetic architectures of individual phenotypes. The results offer novel perspectives on the nature of genetic risk in CHD and the origins of mutational robustness in cardiac development.

Genetic heterogeneity and rare alleles obscure the biological significance of epistasis in human diseases. In contrast, the power to detect genetic interactions in the setting of naturally occurring variation is greater in inbred strain crosses. Through such crosses, we show that the effects of pairwise and higher-order epistasis correlate with the severity of a heart defect in *Nkx2-5*^+/-^ mice. Other features of the genetic architecture, such as the number of QTLs, do not correlate. The QTLs for mild ASDs exert mainly small, additive effects on risk, while the QTLs for severe AVSDs exert large, additive and non-linear effects. The QTLs for moderately severe, ventricular septal defects have intermediate effects. Strikingly non-linear G×G×G_*Nkx*_ effects can either effectively suppress the risk of or account for a large fraction of AVSDs in the *Nkx2-5*^+/-^ population.

A complex genetic model explains the inverse relationship between the severity and incidence of a defect in humans. Whereas a modestly deleterious mutation may be sufficient to cause a mild defect, additional additive or non-linear genetic interactions are necessary to push a mutation carrier past the liability threshold for a severe defect. Severe defects are rarer because the multi-locus genotypes that give rise to deleterious interactions are improbable. Consistent with this model, a small study of non-syndromic AVSD found that among 34 persons who carried a mutation of one of six AVSD genes (*NIPBL*, *CHD7*, *CEP152*, *BMPR1A*, *ZFPM2*, *MDM4*), 8 persons carried two or more, rare or rare, damaging nonsynonymous variants of the six genes.^22^ Larger human studies are necessary to validate an oligogenic model in which non-linear effects may have an outsized role in severe CHD.

The model suggests a corollary: Epistasis should matter less for strongly deleterious mutations that place individuals near the liability threshold for a defect. To exert the same phenotypic effect as a weak mutation and its associated genetic interactions, a strong mutation must either obstruct one critical pathway or perturb several simultaneously. The dramatic effect of *de novo* mutations of histone-modifying genes, which regulate the expression of multiple developmental genes, is consistent with the latter possibility.^23^

The protective and deleterious effects of syn- and anti-homozygosity in G×G×*G*_*Nkx*_ interactions are consistent with genetic coadaptation, a phenomenon first described by Dobzhansky in 1948. In crosses of flies isolated from geographically distant locales, he noted “gene complexes” that conferred greater fitness when their alleles were from the same, rather than different populations.^24, 25^ Increased fitness emerged from the interaction between genotypes rather than their independent effects. Genetic coadaptation is probably widespread but under-recognized. Recent examples relate to growth under different conditions in yeast^8^, male fertility in *Drosophila*,^9^ body composition phenotypes in mice,^26^ and vitamin D receptor and skin color gene coadaptation in humans.^27^ An elegant example examined the genetic regulation of fasting plasma glucose levels, a diabetes-related trait. In a set of mouse crosses between chromosomal substitution strains, one or two different chromosomes from the A/J strain were introduced into the C57BL/6J background. Five interactions were discovered between pairs of A/J chromosomes. Compared to the control, C57BL/6J strain, mice that carried either A/J chromosome of an interacting pair (e.g., chromosome 5 *or* 6) had elevated glucose levels. Under an additive model, mice that carry both A/J chromosomes (e.g., chromosomes 5 *and* 6) should have higher glucose levels, but they actually had normal levels.^28^ A/J and C57BL/6J appear to have arrived at different genetic solutions to maintain normoglycemia.

Finally, the results offer a rare example of selection for genetic robustness.^15^ Whether robustness to mutation arises by selection or as an inherent consequence of an adapted trait has been a longstanding debate.^14, 29^ Depending upon the experimental design, investigators have lacked either the ancestral population to assess the pre-robust state or the years required to detect selection for robustness in an evolving population.^14^ Our application of inbred strains addresses these challenges. Inbreeding strongly selects for fitness under genome-wide homozygosity. Unknown pressures appear to have selected strain-specific genotypes at interacting loci that enhance the robustness of cardiac developmental pathways to subsequent perturbation. *Nkx2-5* mutation was probably not the original selection pressure because there are many potential causes of CHD. Nevertheless, outbreeding in our intercrosses produces individual variation in the robustness to *Nkx2-5* mutation. Modifier locus genotypes permit us to observe the analog of pre- and post-selected states in the same population. If humans similarly vary in the robustness of cardiac developmental pathways, the genetic basis of CHD is a function not only of deleterious variants but also of the degree of coadaptation of genes in the individual.

## METHODS

### Mouse Strains and Crosses

Inbred C57BL/6N (B) and FVB/N (F) mice were purchased from Charles River (Wilmington, Massachusetts) and A/J (A) from the Jackson Laboratory (Bar Harbor, Maine). The *Nkx2-5*^+/-^ mutant line is maintained in the C57BL/6N background.^6, 30^ To produce F_1_ hybrids, *Nkx2-5*^+/-^ C57BL/6N males were crossed to wildtype A/J or FVB/N females. The *Nkx2-5*^+/-^ F_1_ was then crossed to produce A×B and F×B F_2_ progeny. We also produced F10 and F14 advanced intercross generations of F×B by random mating from the F_2_ generation onward. Mating between siblings and first cousins was avoided from the F_3_ and F_4_ generations onward.^31^ Mice were housed under standard conditions in the same room with access to water and chow *ad libitum*. The Washington University School of Medicine Animal Studies Committee approved the experiments.

### Collection and Phenotyping of Hearts

We collected and phenotyped hearts as previously described.^6^ Briefly, pups were collected within hours of birth. Thoraxes were fixed in 10% neutral buffered formalin. After *Nkx2-5* genotyping, all *Nkx2-5*^+/-^ and a subset of wild-type hearts were dissected and embedded in paraffin. Hearts were completely sectioned in the frontal plane at 6 µm thickness. Every section was collected, stained with hematoxylin and eosin, and evaluated by two individuals. Defects were diagnosed by the morphology of the atrial and ventricular septae, semilunar and atrioventricular valves, chambers and the anatomic relationships between them. Secundum ASDs were distinguished from a patent foramen ovale based on the size of the latter in wild-type hearts.

### Single Nucleotide Polymorphism Genotyping

Genomic DNA was isolated by phenol-chloroform extraction. SNP genotyping was performed on affected and normal *Nkx2-5*^+/-^ mice. In the F×B advanced intercross, normal controls were randomly selected from unaffected siblings.

The A×B and F×B F_2_ intercrosses were genotyped using custom panels of ~120 SNPs on the Sequenom MassARRAY system.^32^ The average distance between markers was 19-24 Mb. The F×B advanced intercross generations were genotyped on the Mouse MD Linkage Panel (Illumina, San Diego, CA). A subset of 761 SNPs on the panel were selected for subsequent analyses. We eliminated SNPs that had >10% missing genotypes or low median quality scores (GC < 0.35) or were not in dbSNP (build 141). Fewer SNPs were genotyped in the F2 than the advanced intercross generations because there is less recombination. Combining SNPs from the F×B F_2_ and advanced intercrosses yielded 887 markers spaced an average of 3 Mb.apart. SNP locations were assigned according to GRCm38/mm10. The sex chromosomes were not analyzed. Supplementary Table 2 lists the SNPs.

### Genetic Imputation and Association Analyses

QTLs that modify the risk of a heart defect were mapped via case-control association tests. Each type of heart defect was analyzed separately. The same normal controls were used in each analysis.

We utilized R/qtl (version 1.40-8) to map QTLs under a binary trait model in the A×B F_2_ intercross.^33^ Permutation testing (N = 5000) was performed to determine significance thresholds. Significant and suggestive loci had LOD scores >3.3 (α = 0.05) and >2.7 (α = 0.2) respectively.^34^

We implemented a univariate linear mixed model in GEMMA (version 0.94.1) for a combined analysis of the F×B F_2_ and advanced intercross generations.^35^ A likelihood ratio test compared the alternative hypothesis that a SNP has a non-zero effect against the null hypothesis of zero effect. Each binary phenotype was modeled as a function of SNP genotypes and a random or polygenic effect that accounts for relatedness:

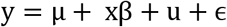

Where y is the vector of binary phenotypes (0 = control, 1 = heart defect), μ is the mean, x is a vector of SNP marker genotypes, β is a vector of marker effects, u is a vector of random effects, and ϵ is a vector of errors. Genome-wide significance thresholds were Bonferroni corrected. The effective number of tests was 542 after accounting for linkage between SNPs.^36, 37^ Significant and suggestive loci had nominal P values < 9.2×10^−4^(0.05/542) and < 1.8×10^−3^(1/542). The combined analysis of the F_2_ and advanced intercross generations enhances the resolution and power of mapping, but poses a couple special considerations.^17^

First, some SNPs were not genotyped in either the F×B F_2_ or advanced intercross generations. They required imputation. In the F_2_, 758 SNPs were imputed using R/qtl.^33^ In the advanced intercross, 126 SNPs were imputed using QTLRel (version 0.2-15). The F_10_ and F_14_ were imputed separately because of differences in recombination patterns between the generations.^38^ We estimated that >80% of missing genotypes in the F_2_ were imputed with >70% confidence; only 2% were imputed with less than 50% confidence. For the advanced intercross generations, 99.9% of missing genotypes were imputed with >70% confidence.

Second, to minimize type I error, we accounted for unequal relatedness between individuals in the advanced intercross generations. The univariate linear mixed model (Equation 1) incorporates a genetic relatedness matrix (GRM) that captures the phenotypic variance due to the genome-wide similarity between individuals. For each case-control analysis, we used GEMMA to estimate centered GRMs using the SNP genotypes from the combined F×B generations. Because F_2_ individuals are essentially full-siblings, their pairwise relatedness coefficients were set to 0.5 and 1 on the diagonal in the GRM. We then adjusted the entire matrix (F_2_ + advanced intercross generations) to the nearest positive definite matrix using the *nearPD* function in the “Matrix” R package (version 1.2-6). The adjustment adds a small amount of noise to the GRM to remove negative eigenvalues, which is necessary for stability of the mixed model. A Mantel test for similarity between the pre- and post-adjusted GRMs showed > 95% similarity in all cases.

### Estimation of Heritability

The proportion of the phenotypic variance explained (PVE) by all genotyped SNPs was estimated using restricted maximum likelihood (REML) in the linear mixed model in GEMMA. This method uses the GRM to estimate the genome-wide PVE and its standard error for each type of heart defect in the A×B and F×B intercrosses.^39^ PVE estimates were corrected for ascertainment bias and transformed to the liability scale (PVE_l_) according to the equation:

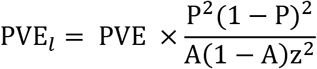

where P is the population disease incidence, A is incidence in the ascertained cohort, and z is the height of the standard normal distribution at the population liability threshold.^40^

### Analysis of G×G_*Nkx*_ Effects

The proportion of the phenotypic variance explained by a SNP, PVE_SNP_, was calculated according to the equation:

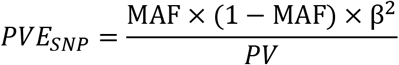

where MAF is the minor allele frequency, PV is the phenotypic variance, β is the effect of the SNP as estimated in GEMMA for both the A×B and F×B intercrosses. PVE_SNP_ estimates were corrected and transformed to the liability scale in the same manner as the genome-wide PVE estimates.

We estimated odds ratios for the G×G_*Nkx*_ effects using logistic regression models. Odds ratios for each A×B locus were obtained using the base glm function in R. Odds ratios for each F×B locus were obtained using the logistic mixed model implemented in GMMAT.^41^ The mixed model controlled for relatedness via the GRMs above.

### Analysis of G×G×G_*Nkx*_ Effects

The contribution of higher-order epistatic effects to the risk of a heart defect was calculated according to the physiological epistasis model.^18^ The G×G×G_*Nkx*_ effect is determined by the difference between the observed and the expected incidences of a defect at a two-locus genotype under the null model of independently acting loci. The expected incidence is calculated from the unweighted regression of the two-locus genotype incidences onto the two single-locus incidences. To compare G×G×G_*Nkx*_ effects between defects that have different incidences, the effects were divided by the unweighted average incidence of the defect across all nine genotype combinations between two loci. Effects that raise or lower risk are deemed positive or negative respectively. To assess statistical significance, we arcsine-transformed the effects to obtain unbiased variances.^42^ Using these variances, we conducted two-tailed T-tests to identify significant, non-zero effects. The significance threshold was Bonferroni-corrected for the number of two-locus genotypes in a pairwise analysis, i.e., P = 0.05/(9×), where x is the number of pairwise combinations of loci for a defect. To compare effects between defects that have different incidences, the significant epistatic effects were divided by the unweighted average incidence of each defect.^18^ Because only one G×G_*Nkx*_ modifier locus was identified for ASD and muscular VSD in the F×B population, we assessed G×G×G_*Nkx*_ effects in combination with the next most significant SNP (ASD: rs13478997, Chr. 6, P = 1.94×10^−3^; muscular VSD: gnf13.115.241, Chr. 13, P =2.39×10^−3^.)

### Statistical Analyses

GEMMA was run on the Ubuntu operating system (version 14.04 LTS). All other statistical analyses were performed in the R Statistical Computing environment (version 3.3.1). All t- and z-tests for comparisons of means and proportions respectively were two-sided. We used the R package “ppcor”^43^ to calculate Kendall’s (non-parametric) partial rank correlation τ with a two-sided alternative hypothesis. Partial correlations permitted us to calculate non-parametric correlation while controlling for the intercross and the sample size.^44, 45^ For the partial correlation calculation of G×G×G_*Nkx*_ effects versus defect severity (Fig. 3e), we controlled for variation in the observed incidence at two-locus genotypes to avoid bias by an underlying correlation.

## SUPPLEMENTARY MATERIALS

Supplementary Tables 1 and 2 and Supplemental Figures 1-3 are included in supplementary materials.

## Acknowledgements

Author contributions: E.A. and P.Y.J. conceived the analyses and project. E.A., S.D.R., R.A.M., N.U., Y.Q., C.E.S. and P.Y.J. performed experiments. E.A. performed statistical analyses. E.A., J.M.C., and P.Y.J. analyzed data. E.A. and P.Y.J. wrote the manuscript with input from J.M.C. and review by all other authors. P.Y.J. supervised the project. We thank Shin-ichiro Imai, Kenneth Olsen, Christoph Preuss, Alan Schwartz, Alan Templeton, Anna L. Tyler and David Wilson for their feedback. Xiang Zhou provided helpful guidance with relatedness matrices in GEMMA.

