## Supplementary Figures 1-3 for "The fitness cost of a congenital heart defect shapes its genetic architecture"

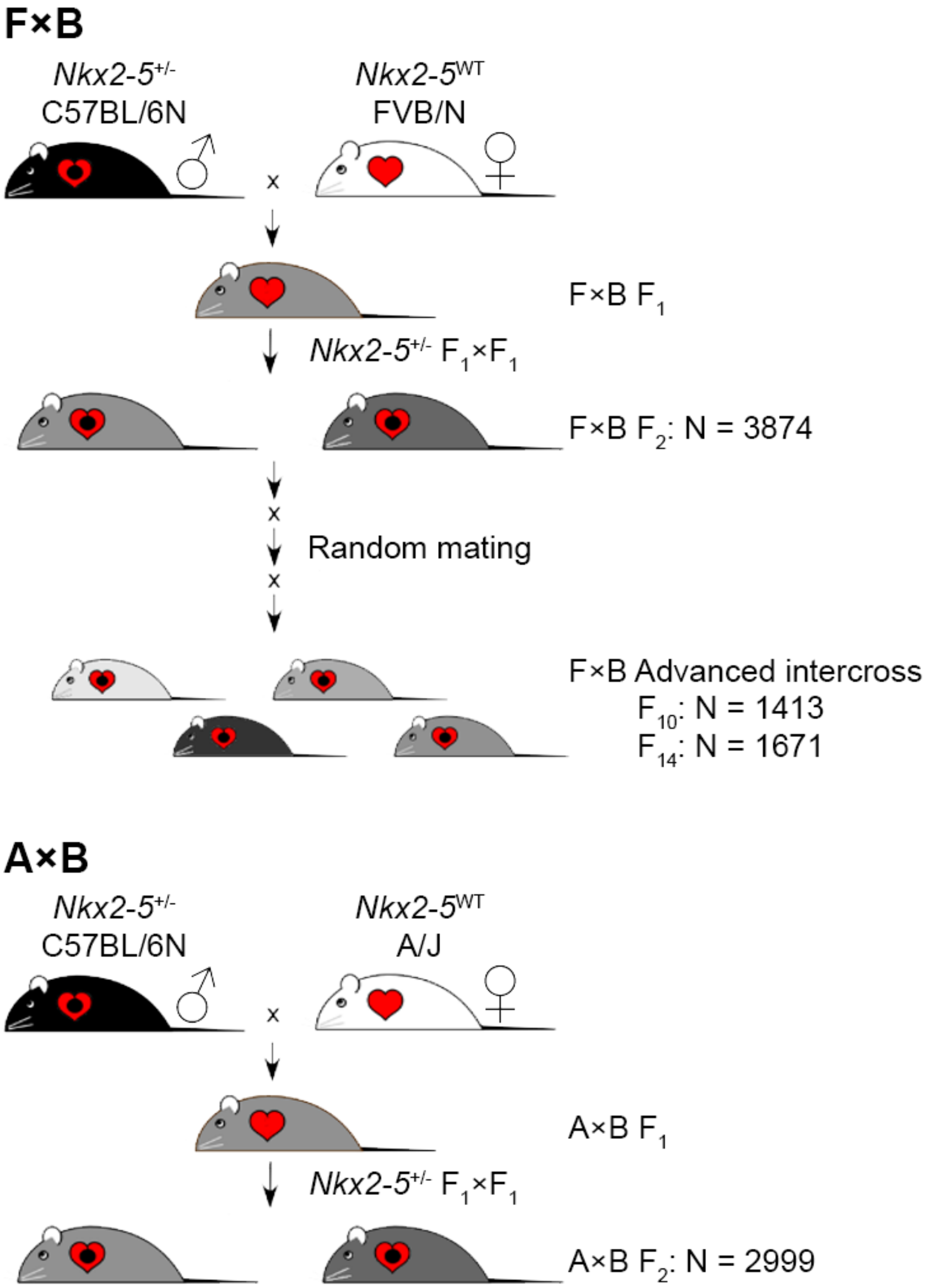


**Supplemental Figure 1. Breeding scheme for inbred strain intercrosses.** We produced $F_1$ hybrids by crossing wild-type FVB/N (F) or A/J (A) females to *Nkx2-5*$^{+/-}$ C57BL/6N (B) males. We phenotyped newborn *Nkx2-5*$^{+/-}$ $F_2$ pups from both A×B and F×B intercrosses and $F_{10}$ and $F_{14}$ pups from the F×B advanced intercross. The advanced intercross was produced by random mating of non-siblings and non-first cousins beginning in the $F_3$ and $F_4$ generations. The number of *Nkx2-5*$^{+/-}$ pups phenotyped in each generation is shown.

**a**

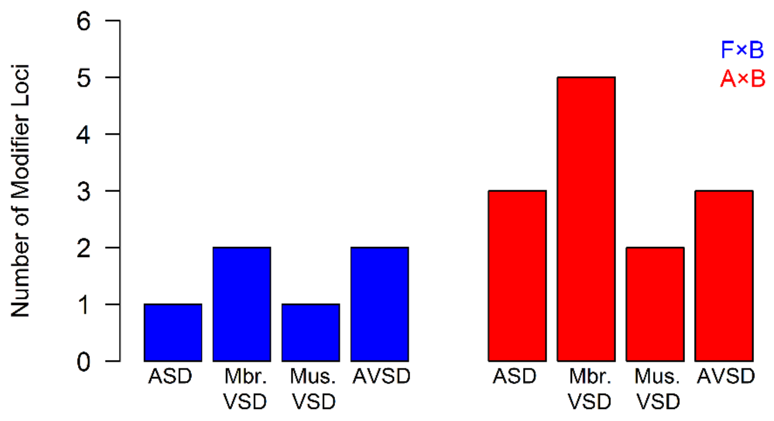


**b**

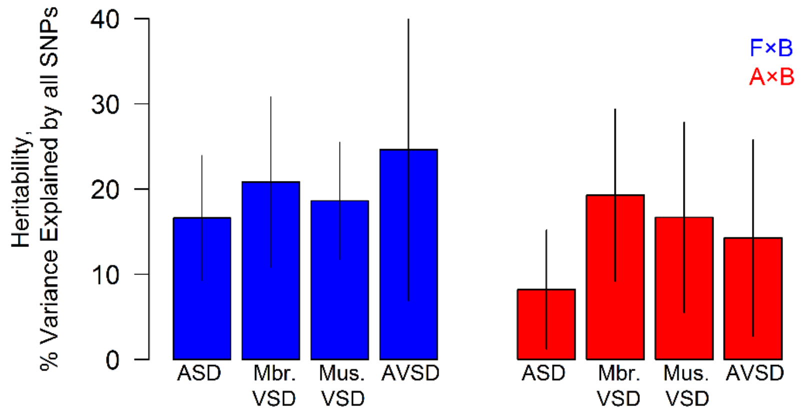


**Supplemental Figure 2.** Neither **a,** the number of significant and suggestive loci detected nor **b,** the heritability explained by all genotyped SNPs varies with the severity of a defect. Error bars are 95% C.I.

**Supplemental Figure 3.** The observed incidences at two-locus genotypes of ASD, membranous and muscular VSD, and AVSD are shown between every pair of suggestive or genome-wide significant modifier loci found in both crosses. In the two cases where there is only one suggestive locus, the locus is plotted against the next most significant locus in the cross.

**ASD: F×B**

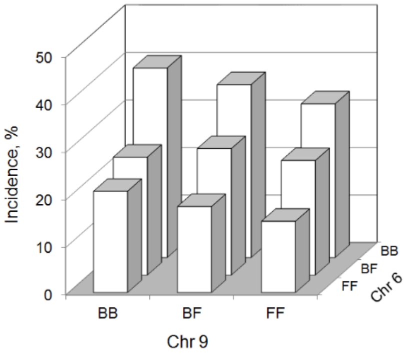


**ASD: A×B**

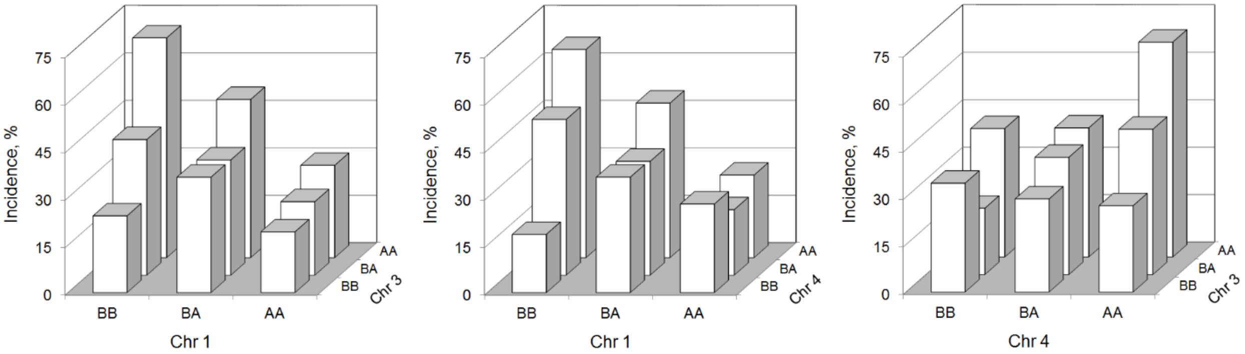


**Membranous VSD: F×B**

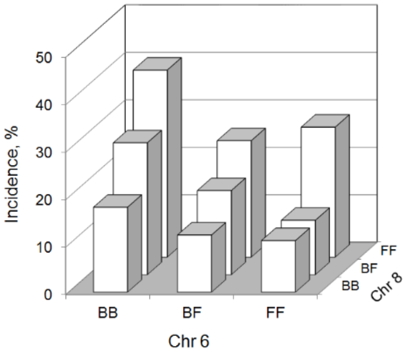

### Membranous VSD: A×B

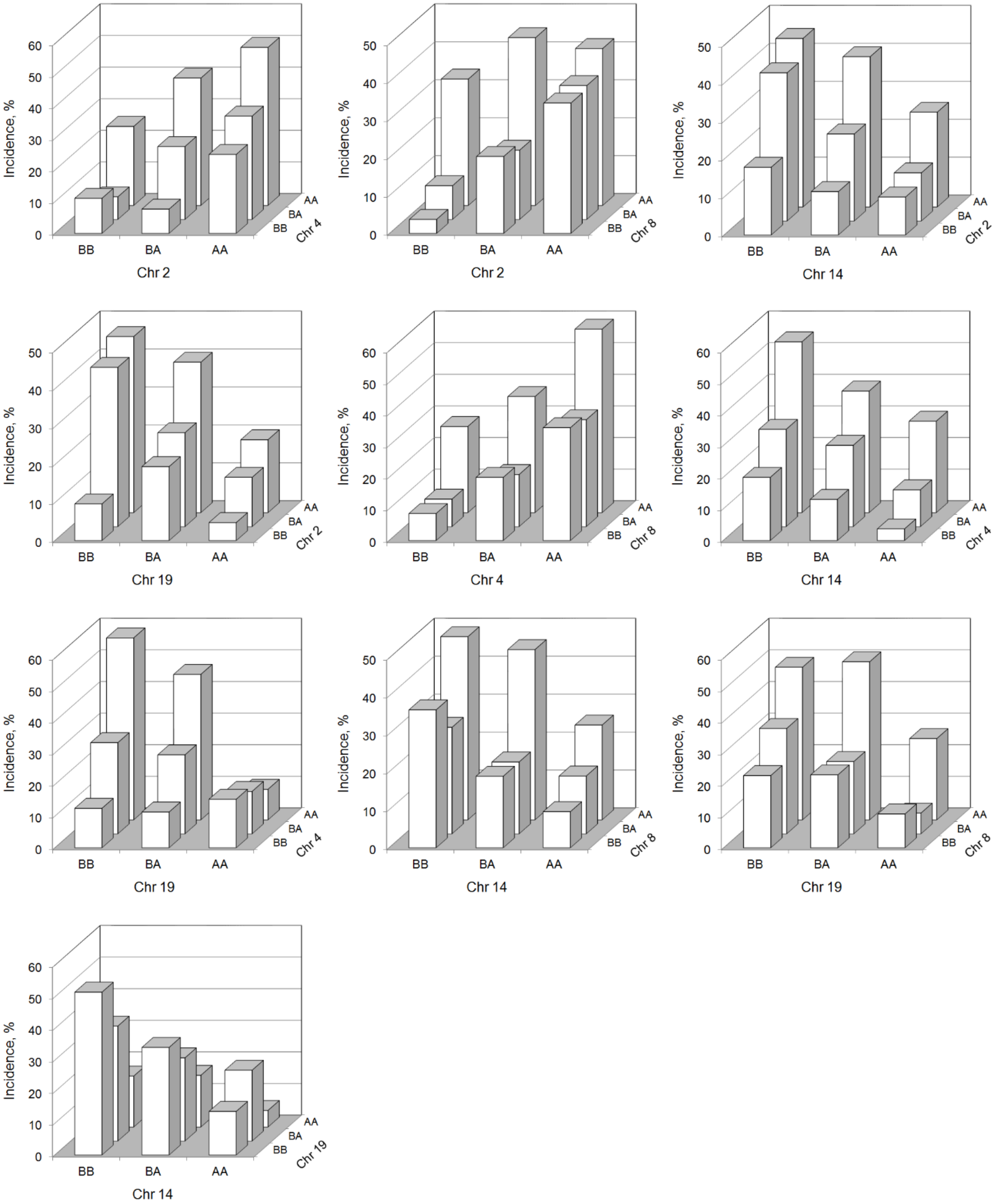

### Muscular VSD: F×B

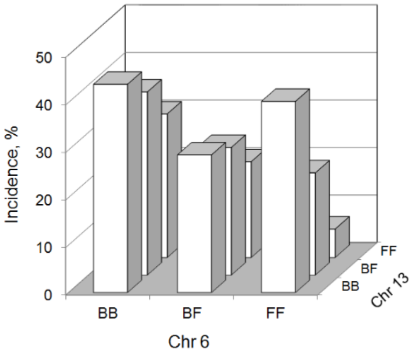


### Muscular VSD: A×B

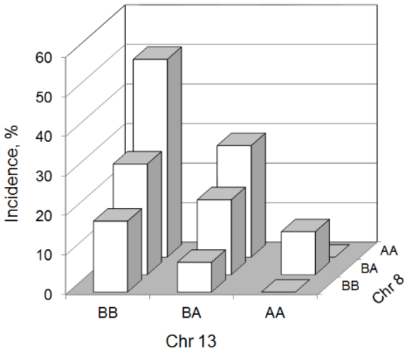

### AVSD: F×B

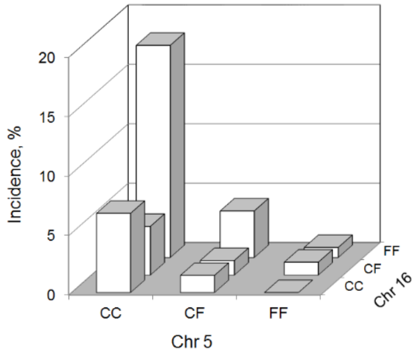


### AVSD: A×B

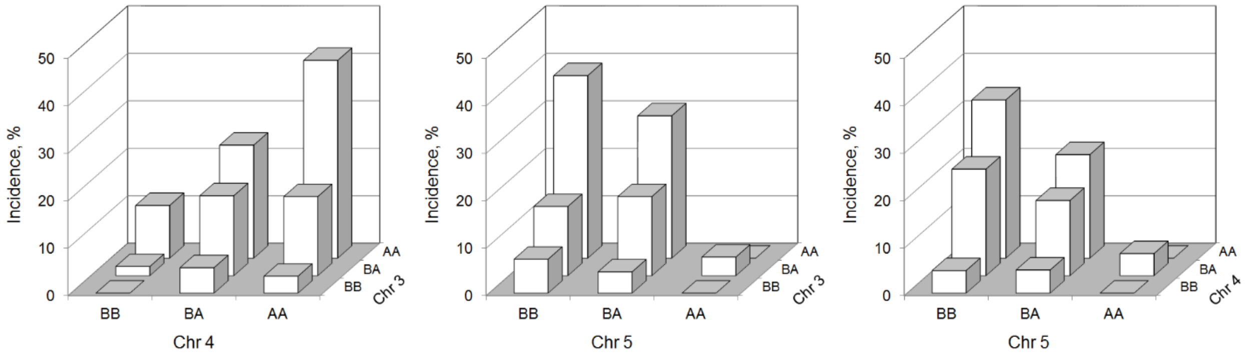
